# Trait differentiation across urbanized environments in invasive *Centaurea melitensis* is not predicted by the Plant Economic Spectrum

**DOI:** 10.64898/2026.10.30.762855

**Authors:** Anthony J. Dant, Lily G. Bishop, Katrina M. Dlugosch

**Affiliations:** All University of Arizona, Tucson, AZ, 85721

**Keywords:** Centaurea melitensis, California, Urban Ecology, Plant Economic Spectrum, Species Invasion

## Abstract

Urbanization imposes unique selective pressures on organisms, but connecting specific urban environmental differences to trait adaptation requires detailed characterization of environmental heterogeneity across urbanized landscapes and tests for adaptation to it, rarely done to date. The Plant Economic Spectrum (PES) predicts that plants invest in different life strategies according to resource availability, ranging from an acquisitive strategy favoring fast growth and aboveground biomass where resources are abundant, to a conservative strategy favoring belowground investment where resources are limiting. Here, we leverage previous work describing the mosaic of urbanized environments in California, USA as a set of Environmental Zones, and use this framework to test for trait differentiation predicted by the PES in the invasive annual plant *Centaurea melitensis*. Seeds from 31 populations across California were grown in greenhouse conditions for two generations. In the second generation, we measured growth, reproductive, and phenology traits to test for genetically-based differentiation and whether variation follows PES predictions. Variation across all genotypes was not structured along the multivariate trait axes predicted by the PES. Individual traits were associated with environmental variables, including impervious surface cover and tree cover, following some PES expectations, but multi-trait strategies were absent. We found evidence that *C. melitensis* traits have differentiated genetically across environments with different features of urbanization but did not exhibit the multi-trait strategies predicted by the PES. Our study pairs detailed knowledge of urbanized environments with tests of trait-environment relationships to reveal how urbanization may be selecting on the species that inhabit these spaces.

## INTRODUCTION

Urban environments are now recognized as unique habitats that can select for adaptations in the organisms that live there (Johnson & Munshi-South, 2017; Alberti et al., 2020; Johnson et al., 2024). A growing number of studies have found evidence for changes in behavior, morphology, and physiology associated with urban environments (Hendry et al., 2008; Isaksson, 2020; Ritzel & Gallo, 2020; Corsini et al., 2021; Santangelo et al., 2022). However, in most cases neither the specific aspects of an urbanized environment that select on an organism, nor how these are associated with specific adaptations, are yet understood (Rivkin et al., 2019; Miles et al., 2021; Leandro et al., 2025). Building this understanding requires at least two things 1) a detailed characterization of what differs in an urbanized environment (Donihue & Lambert, 2015; Lambert et al., 2021; Szulkin et al., 2020), including not only climate but other unique factors affecting organisms as well, such as patterns of pollution and poverty (Leong et al., 2018; Schell et al., 2020; Des Roches et al., 2021; Johnson et al., 2024), and 2) tests of hypotheses about how these environmental differences select on traits (Ramalho & Hobbs, 2012; Diamond et al., 2018; Campbell-Staton et al., 2020; Diamond & Martin, 2020; Miles et al., 2021; Thompson et al., 2022; Verrelli et al., 2022).

Regarding the first step of identifying how the environment differs in urbanized areas, here we leverage a study system in which the complex environmental variation across an urbanized landscape has been exceptionally well-characterized. *Centaurea melitensis* (L., Asteraceae, ‘Maltese starthistle’) is an annual forb, native to Northern Africa, Southern Europe and Western Asia (DiTomaso and Healy 2007). It has been introduced to every continent except Antarctica (Hellwig 2004). It predominantly inhabits grassland, oak woodland, and chaparral environments (DiTomaso and Healy 2007). It has also established in a variety of urban habitats (Roché and Roché 2021; Dant et al. 2025). In North America, the oldest record of *C. melitensis* occurs in southern California in 1789 (Hendry 1931), suggesting the species has been colonizing the region for over 200 generations. It easily self-fertilizes via flowering heads that do not open (cleistogamous capitula), but also has the ability to reproduce by open flowerheads, with unknown potential to outcross (Porras and Muñoz 2000). There is previous evidence for both local adaptation and plasticity across its populations, in response to nitrogen availability and aridity (Moroney et al., 2013; Humphries et al., 2024; Valliere et al., 2025), suggesting its potential to adapt to local urban environments.

We have previously characterized environmental variation across the mosaic of urbanized landscapes occupied by *C. melitensis* in California, USA (Dant et al. 2025; Appendix A of this dissertation). California includes six large cities with populations over 500,000, many smaller cities, numerous natural areas that border metropolitan and agricultural development, and a wide range of socio-economic, climatic, and environmental conditions (Berg et al., 2024). California also organizes spatial and environmental data in easily accessible, publicly available databases, such as those provided by the California Environmental Protection Agency (CALEPA; https://calepa.ca.gov), and the Multi-Resolution Land Characteristics Consortium (MRLC; https://www.mrlc.gov). Our previous study utilized a hierarchical cluster analysis to classify environmental variation across California at the census tract level, based on 19 environmental variables across 7,829 census tracts (Dant et al. 2025). Environmental variables included land cover, climate, pollution, and socio-economic attributes. This analysis resulted in the identification of nine ‘Environmental Zones’ capturing environmental variation across the mosaic of urbanized and natural environments in California. An initial analysis of plants from wild collected seed from 17 populations of *C. melitensis* revealed trait differences among populations in different Environmental Zones (Dant et al. 2025).

This foundation of knowledge about environmental and trait variation among *C. melitensis* populations in California makes this system highly suitable for testing hypotheses about how urban environments might select on traits. In plants, a core hypothesis connecting the environment to adaptations in traits is the Plant Economic Spectrum hypothesis (PES; Reich, 2014). Building on the Leaf Economic Spectrum hypothesis (Reich et al., 1999; Wright et al., 2004), the PES postulates that across species, plant traits tend to vary together along a continuum, from a resource-conservative strategy that is adaptive in in stressful, resource-limited environments, to a resource-acquisitive strategy that is adaptive in less stressful, resource-abundant environments (Reich, 2014) (Figure 1). For instance, the PES predicts that resource-conservative plant species will be found in more arid environments, where temperature is higher and water more limited; they should have higher root biomass, smaller and longer-lived leaves, and shorter stems than species in cooler and wetter environments (Reich, 2014). The PES has become a core hypothesis for predicting the life history strategies of plants in ecology, with the foundational paper by Wright et al. 2004 cited over 11,000 times as of this writing (Google Scholar, accessed August 12th 2026).

**Figure 1.**
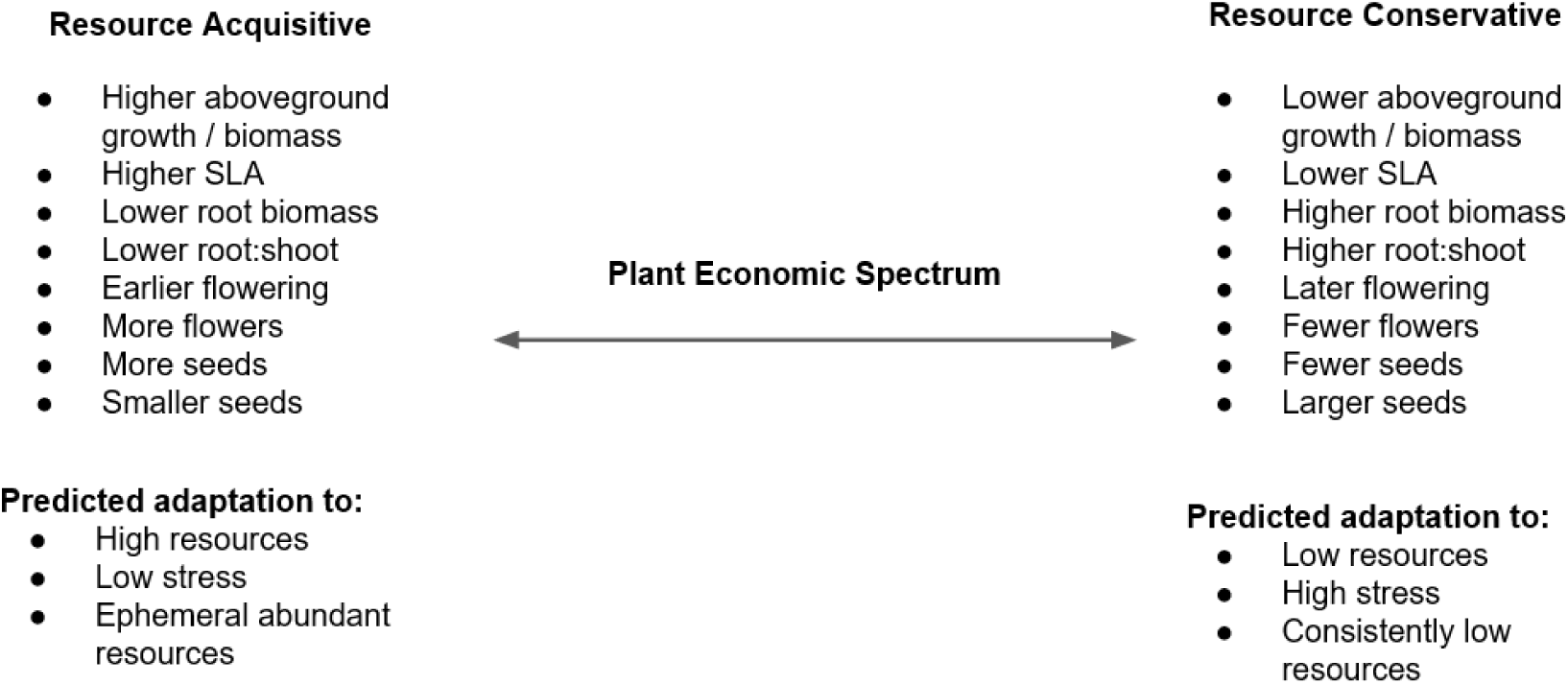
Conceptual diagram of trait differences as predicted by the Plant Economic Spectrum (PES), based on (Reich 2014), for traits measured in this study. Urban environments may include more stressful habitats that select for a Resource Conservative strategy, or ephemeral abundant resources that select for a Resource Acquisitive strategy, but the existence of a PES in annual plants has been debated (Brouillette et al., 2014; Kurze et al., 2021).

Though the PES has been primarily used to describe adaptive life history strategies across perennial species (Freschet et al., 2010; de la Riva et al., 2016, Montesinos, 2022), there has been growing interest in applying this influential framework to annual species as well, and to using it to explain differences among populations of a single species (Brouillette et al., 2014; Kurze et al., 2021). Urban environments in particular often generate a ‘heat island’ effect that results in elevated temperatures. This effect has been associated with changes in plant traits (Calfapietra et al., 2015; Johnson et al., 2015; Čeplová et al., 2017). The PES would predict that hotter urban environments would select for traits typical of a more resource-conservative strategy (Figure 1). Alternatively, urban environments that provide abundant but ephemeral resources would be predicted to select for a resource acquisitive strategy (Figure 1). Testing these predictions requires a nuanced understanding of what characterizes different urban environments, which is missing in most systems, and to our knowledge these predictions have not yet been tested.

Here we use a sample of 31 populations of *C. melitensis*, reared for two generations in a common environment, to test whether genetically-based trait variation in this system follows the predictions of the PES (Figure 1). First, we ask whether variation across all genotypes from all populations is structured along the trait axes predicted by the PES. Next, we test for trait differentiation across our previously-identified Environmental Zones that describe urbanized landscapes in California (Figure 2). After identifying significant trait differentiation across Zones, we test for trait correlations with specific environmental variables that characterize those Zones. Finally, we ask whether patterns of trait differentiation across environments align with predicted trait-environment relationships from the PES (Figure 1). Our study brings together detailed knowledge of complex urbanized environments with tests of hypothesized trait-environment relationships to gain insights into how urbanization might be selecting on the species that inhabit these spaces.

**Figure 2.**
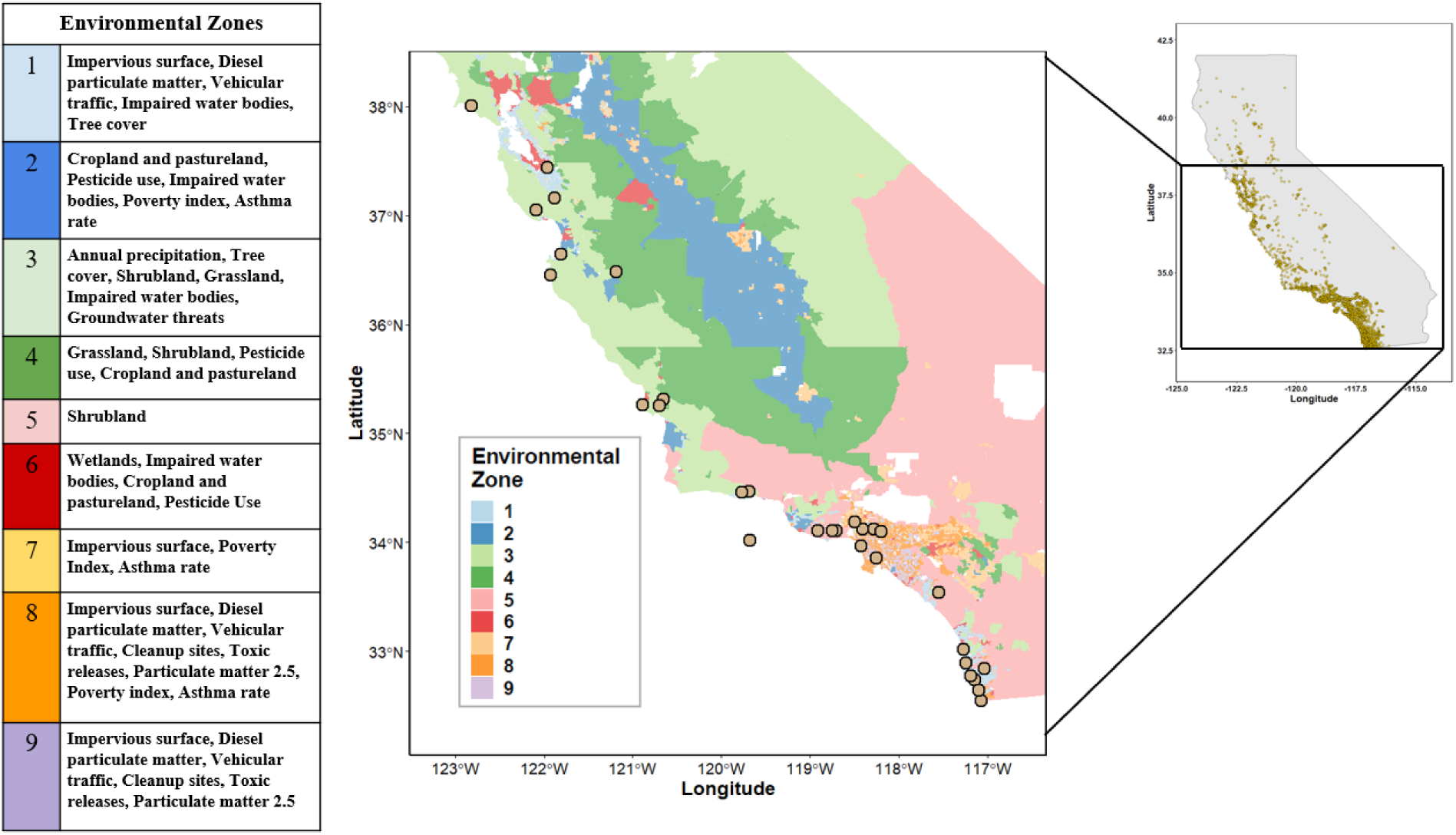
Map of collection sites in the State of California (center) colored by the nine Environmental Zones (left) as defined by Dant et al. (2025). Circles indicate the 31 collection sites that were included in this study, including Santa Cruz Island in the Pacific Ocean. Dominant environmental characteristics of each Environmental Zones are noted at left. Righthand map shows the current *Centaurea melitensis* range in California (yellow dots) obtained from the Global Biodiversity Information Facility (GBIF.org (24 June 2026) GBIF Occurrence Download https://doi.org/10.15468/dl.48njm5).

## MATERIALS AND METHODS

### Collections

Seeds from 31 *C. melitensis* populations were collected across California from 2021 to 2022. Sampling locations are illustrated in Figure 2. These collection sites included multiple areas of major cities, such as Los Angeles and San Diego, as well as surrounding areas, across a range of Environmental Zones defined by Dant et al. (2025) (Figure 2). Seeds were collected from individual plants (maternal lineages) within each site, located at least 1m apart along a linear transect. The total number of lineages collected and coordinates for each population can be found in the supplemental materials (Table S1).

### Greenhouse experiment

We grew *C. melitensis* for two generations within a greenhouse environment to minimize maternal/environmental effects on seeds and test for genetically-based differences in traits among populations. Field collected seeds were grown in a common environment and their progeny from self-fertilization were collected for use in the greenhouse experiment. To do this, seeds from each lineage at each site were germinated in a greenhouse at the University of Arizona (Tucson, AZ, USA) and grown to sexual maturity. Flowering heads were covered with white organza bags (TheDisplayGuys, Ontario, CA, USA) and allowed to self-fertilize. Seeds from this second generation were collected and placed into petri dishes containing commercial potting soil (Happy Frog Potting Soil, FoxFarm Soil and Fertilizer Co, Arcata, CA, USA) under 12 h of fluorescent lights at 21 °C until they germinated. Individual seedlings were then transplanted into 410 ml Deepots (Steuwe & Sons Inc, Tangent, Oregon, USA) containing one of two substrate combinations: A) a typical ‘soil’ treatment consisting of 1 part sand (30 grit Quikrete, The QUIKRETE Companies, Atlanta, GA, USA) to 5 parts soil (Happy Frog Potting Soil) / coconut husk mixture (Chips-N-Fiber Premium Coconut Husk, PROCO, Tucson, Arizona, USA); or B) a high ‘sand’ treatment to allow the harvesting of roots, consisting of 1 part sand to 1 part soil / coconut husk mixture. Plants in the soil treatment had no destructive growth measurements taken and were grown to maturity, while plants in the sand treatment were harvested earlier and were used for all destructive trait measurements (see below). Plants were grown from August 2023 to April 2024 under ambient light and 24 °C average temperature in the greenhouse. Final sample sizes for plants available for all measurements were 7 to 24 from each of the 31 sites in the soil treatment, and 7 to 16 per site in the sand treatment. Within treatments, each plant represented a different maternal lineage from each site.

Two weeks after transplanting, growth trait measurements began. Growth traits included (a) relative growth rate (RGR) of the length and width of the longest leaf, and RGR of the total number of leaves; (b) Specific leaf area (SLA); (c) Root diameter; (d) Total root and aboveground biomass; and (e) Root to shoot ratio. RGR of the length and width of the longest leaf, as well as the number of leaves, were calculated from two measurements, the first at two weeks after transplanting and the second after an additional two weeks.

SLA, root diameter, total root biomass, total aboveground biomass, and root to shoot ratio were measured prior to bolting for individuals within the sand treatment. SLA was determined for a single leaf per individual by measuring leaf areas with the Leafscan app (Anderson and Rosas-Anderson 2017 Leafscan Version 1.3.21 (retrieved from https://itunes.apple.com/app/id1254892230) followed by drying using silica gel for at least three days for dry mass measurements. Root diameter was measured with calipers. Biomass was measured after oven drying at 65 °C.

Four reproductive trait measurements were taken for each plant in the soil treatment: the date of the first flower, total number of flowerheads (capitula), number of seeds produced by each flowerhead, and average seed weight from five seeds per individual. The total number of flowerheads and seeds were measured at the time of senescence.

### Statistical analyses

To identify whether our lineages, across all sites, harbored variation in PES traits (as predicted in Figure 1), along which they might have evolved differences between sites, we conducted a Principal Components Analysis (PCA). First, pairs of traits that were highly correlated across all lineages from all sites, and therefore redundant for further analyses, were identified using a Pearson correlation matrix among all trait variables, using ggcorrplot v0.1.4.1. in R (v4.5.0 2025) via RStudio version 4.1.3 (RStudio Team 2023). Any pair of traits that exceeded a correlation of r > |0.7| were identified and only one of the pair was retained. A PCA was then constructed to identify axes of trait variation among the retained traits using factoextra (Kassambara & Mundt, 2020) in R. We compared these axes of trait variation across all samples to expected axes of variation of the PES.

Due to a lack of strong axes of multivariate trait variation in the PCA (see Results), we tested for evolved differences across environments for all traits individually. Linear mixed models were used to test for trait differentiation with fixed effects of environment, and significant trait differentiation across environments was compared to predictions from the PES (Figure 1). First, traits were modeled with fixed effects of the Environmental Zone they were assigned from Dant et al. 2025 (two sites were not classified in that study and were assigned the environmental zone of the closest census tract; Table S1), with additional fixed effects of collection location nested within Environmental Zone, and of block position in the greenhouse treated as a random effect (where blocks in this greenhouse are known to differ slightly in environment according to their proximity to cooling units in this greenhouse; Cang et al. 2024). A Bonferroni correction was applied when assessing the significance of each, to account for multiple tests of the same hypothesis of trait differentiation (Dunn 1961). For models with significant trait differences among Environmental Zones, Sidak’s post hoc tests were used to identify individual Environmental Zones with significantly different traits, using emmeans v2.0.1 in R (v4.5.0 2025) via RStudio version 4.1.3 (RStudio Team 2023).

Once Zones with significantly different traits were identified, linear mixed models were used to model traits with fixed effects of specific environmental variables associated with those zones, with block included as a random effect. All traits were included in these analyses (not just those that had significant differences among Zones) to identify potential multi-trait differences across environments, as predicted by the PES (Figure 1). Again Bonferroni corrections were applied to account for repeated tests across traits. All linear mixed models used the lme4 package (Bates et al. 2015) in R.

Specifically, our analyses indicated that multiple traits were significantly different in populations from Environmental Zone 4 relative to the other Zones (see Results). Environmental Zone 4 is dominated by grassland, shrubland, crop, and pastureland, with a high level of pesticide use (described in Figure 2, Figure S1). According to details of variables dominating each Zone from Dant et al. (2025) (provided in Figure S1), we identified three variables that differentiated Environmental Zone 4 from most others:

- <u>Pesticide use</u>: Pesticides are a direct human impact on the environment and are higher in Environmental Zone 4 than others occupied by *C. melitensis*. They are associated with agricultural environments in this Zone, which are typically thought to provide abundant but ephemeral resources for weedy species such as *C. melitensis* (Petit et al., 2011; Chauhan et al., 2012). This would predict a more resource-acquisitive strategy in areas of high pesticide use relative to areas with lower use. Pesticide percentile for each site (percentile of the total pounds of active pesticide ingredients used in agriculture per square mile across the state of California) was obtained from the CALENVIRO 4.0, 2023 report (accessed on April 2025).
- <u>Impervious surface area</u>: Impervious surfaces (human-made structures) are another direct human impact on the environment and are lower in Zone 4 than other zones occupied by *C. melitensis*. Impervious surfaces are associated with heat islands and these are predicted to lead to shorter growing seasons available under the higher temperatures (Johnson et al., 2015; Čeplová et al., 2017; Woudstra et al., 2024). A shorter available growing season would predict a more resource-aquisitive strategy in areas with higher impervious surfaces relative to areas with lower impervious surface area. Impervious surface area percentage was obtained from the MRLC (https://www.mrlc.gov; accessed on April 2025), calculated within a 30 meter boundary of each collection site using the Zonal Statistic tool in ArcGIS Pro (v3.03) as in Dant et al. (2025).
- <u>Tree cover</u>: Tree cover was lower in Environmental Zone 4 than other zones occupied by *C. melitensis*. Tree cover in California increases with higher water availability, and it is made up of a combination of forested areas and suburban human landscaping in areas that would otherwise be grass or shrublands (Clarke et al., 2013; Avolio et al., 2015). Higher water availability would predict a more resource-acquisitive strategy in areas with higher tree cover relative to areas with lower tree cover. Tree cover percentage was obtained using the same methods as for impervious surface area.

## RESULTS

### Trait variation

We retained nine of the twelve traits measured for analyses. The total number of seeds and flowerheads per individual were strongly positively correlated (r = 0.77; Figure S2). Some plants lacked seed count data due to missing seed heads, and therefore we retained flowerhead number and removed seed number from further analyses. Additionally, total aboveground biomass, total root biomass, and root diameter were strongly, positively correlated with one another (all correlations r > 0.7). Among these three variables, we retained only aboveground biomass for downstream analyses. Moderately positive correlations were also present among traits measured on the same tissues, including the RGR of leaf width and length (r = 0.56), RGR of leaf length and number (r = 0.48), and root mass and root:shoot ratio (r = 0.49). All of these variables were retained due to their partial independence. Finally, there was one moderately negative correlation between the number of flowers produced and the date of the first flower (r = -0.45), such that plants that produced flowers earlier tended to produce more flowers, and again we retained both traits.

All traits harbored variation among lineages, but the PCA of the nine retained traits did not show the trait correlations expected under the PES (Figure 1). Across all lineages from all sites, the PCA did not show negative correlations between traits associated with resource acquisition versus those associated with allocation strategies, as would be expected under a fundamental multivariate trade-off (Figure 3). Moreover, PC axes 1–4 explained relatively low fractions of variation on each axis (25.2%, 16.9%, 13.7%, and 10.9% of the variation, respectively), indicating that traits did not covary (as would be expected for multivariate trait strategies) and several dimensions were required to describe variation in traits. Most traits were strongly correlated with both the first and second axes (shown by long vectors), and not closely aligned in direction with either, indicating a lack of a dominant axis of plant strategies across these lineages. PC1 reflected positive correlations among nearly all traits, such that the axis varied from large to small plants (both above and belowground), with only date of first flower varying in the opposite direction, such that earlier flowering was associated with larger and more fecund plants. PC3 was dominated by SLA and root:shoot ratio, which were positively correlated with each other along this axis, opposite of predictions from the PES (Figure 1). PC4 was dominated by the single trait of seed mass.

**Figure 3.**
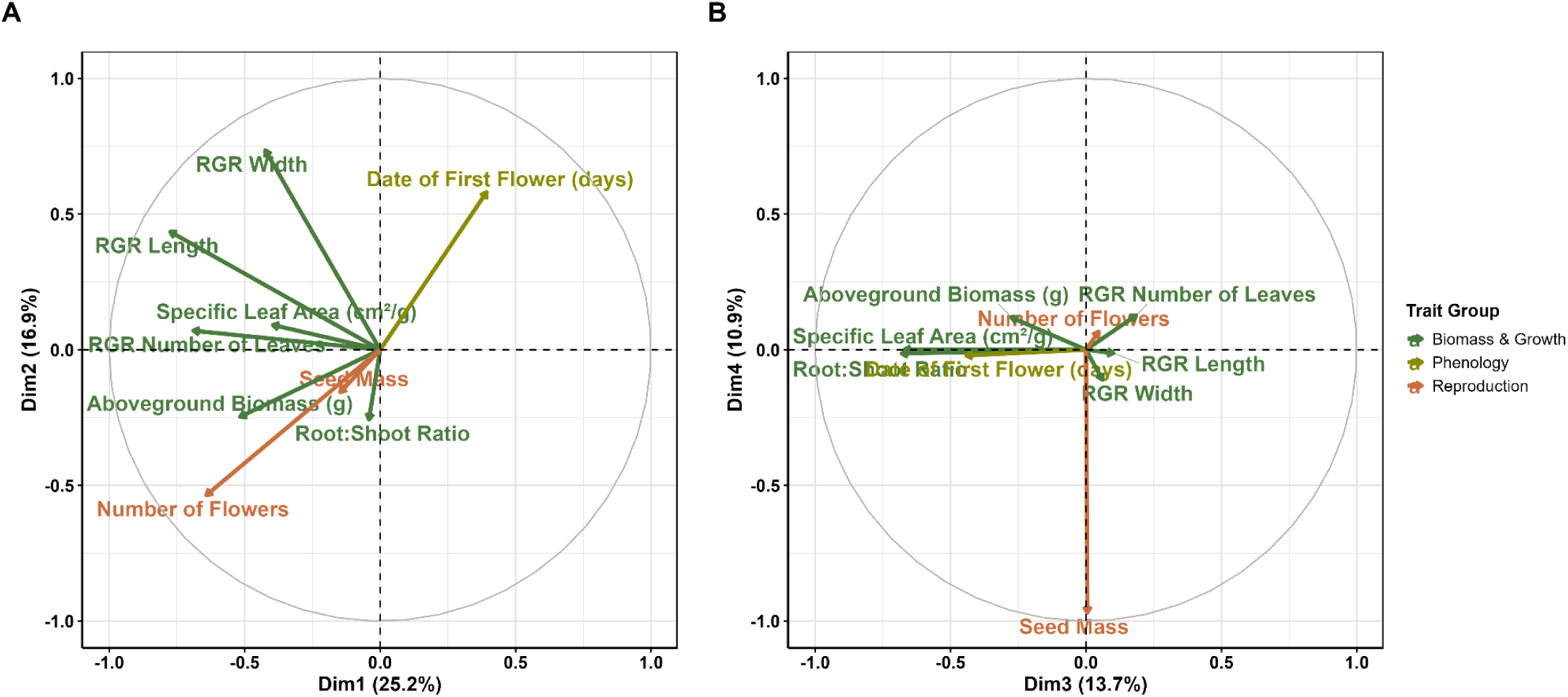
Principal Components Analysis of nine traits measured in the greenhouse experiments including biomass (green), phenology (yellow), and reproductive (orange) traits. Shown are a) PC axes 1 and 2 and b) PC axes 3 and 4, with the percent of variation explained for each indicated in parentheses. Vectors show how traits (labeled) contribute to each axis (vector direction) and the strength of their correlation with these dimensions (vector length).

### Trait differentiation among environments

Our models indicated that there were significant, genetically-based differences in traits of lineages from different environments, identified both when using Environmental Zones and when using individual variables to characterize environments. When modeling traits with fixed effects of the Environmental Zones from which lineages were derived (Figure 2), two traits (RGR of the length of the longest leaf and RGR of the number of leaves) were statistically higher in lineages from Environmental Zone 4 relative to most other zones (Figure 4; model statistics in Table S2). In addition, the date of first flower was significantly higher in lineages from Environmental Zone 3 than Zone 6 (Figure 4; Table S2). Finally, the model for RGR of the width of the longest leaf showed a significant effect of Environmental Zone before Bonferroni correction but not after, and there were no significant pairwise differences in Sidak post hoc tests (Figure 4; Table S2).

**Figure 4.**
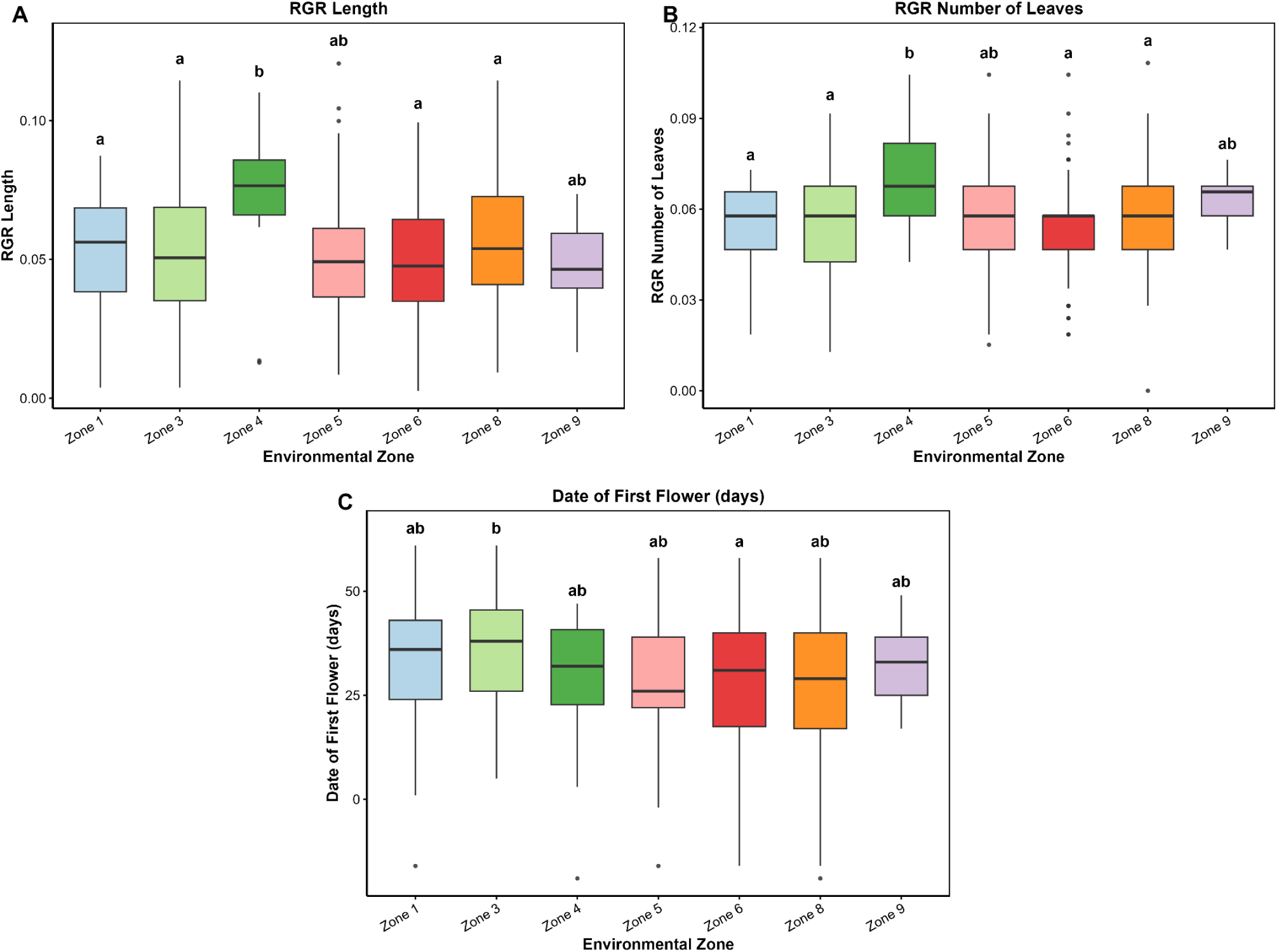
Significant trait differentiation of *Centaurea melitensis* across Environmental Zones for four traits A) relative growth rate (RGR) of the length of the longest leaf, B) RGR of the number of leaves C) date of the first flower. Letters above the box plots indicate significantly different groups as determined by post hoc tests of linear models.

Based on these results, we further tested how trait differentiation was explained by three environmental variables that differed between Environmental Zone 4 and most other Zones (see Methods). Linear mixed models identified four associations between measured traits and these environmental variables that were significant before Bonferroni correction, and one that was significant after(Figure 5; Table S3):

- Pesticide use:

- RGR of the number of leaves was positively related to pesticide percentile, but was not significant after correction (F = 4.133, p = 0.0426). This relationship itself is consistent with what would be predicted by a more resource-acquisitive strategy in areas with higher pesticide use, but no other traits had significant relationships with pesticide use.
- Impervious surface area:

- The date of the first flower was negatively related to impervious surface area, such that lineages from areas with higher impervious surface cover flowered earlier, but this was not significant after correction (F = 4.666, p = 0.0314). This relationship is consistent with what would be predicted by a more resource-acquisitive strategy in areas with higher impervious surface area, but no other traits had significant relationships with impervious surface area.
- Tree cover:

- RGR of number of leaves was negatively related to tree cover, and was significant after correction (F = 9.325, p = 0.0024). This relationship is opposite of what would be predicted for a more resource-acquisitive strategy with more tree cover.
- The root-to-shoot ratio was positively related to tree cover, such that lineages with more root investment were from areas with more tree cover, but this was not significant after correction (F = 5.486, p = 0.0196). This relationship is opposite of what would be predicted for a more resource-acquisitive strategy with more tree cover.

**Figure 5.**
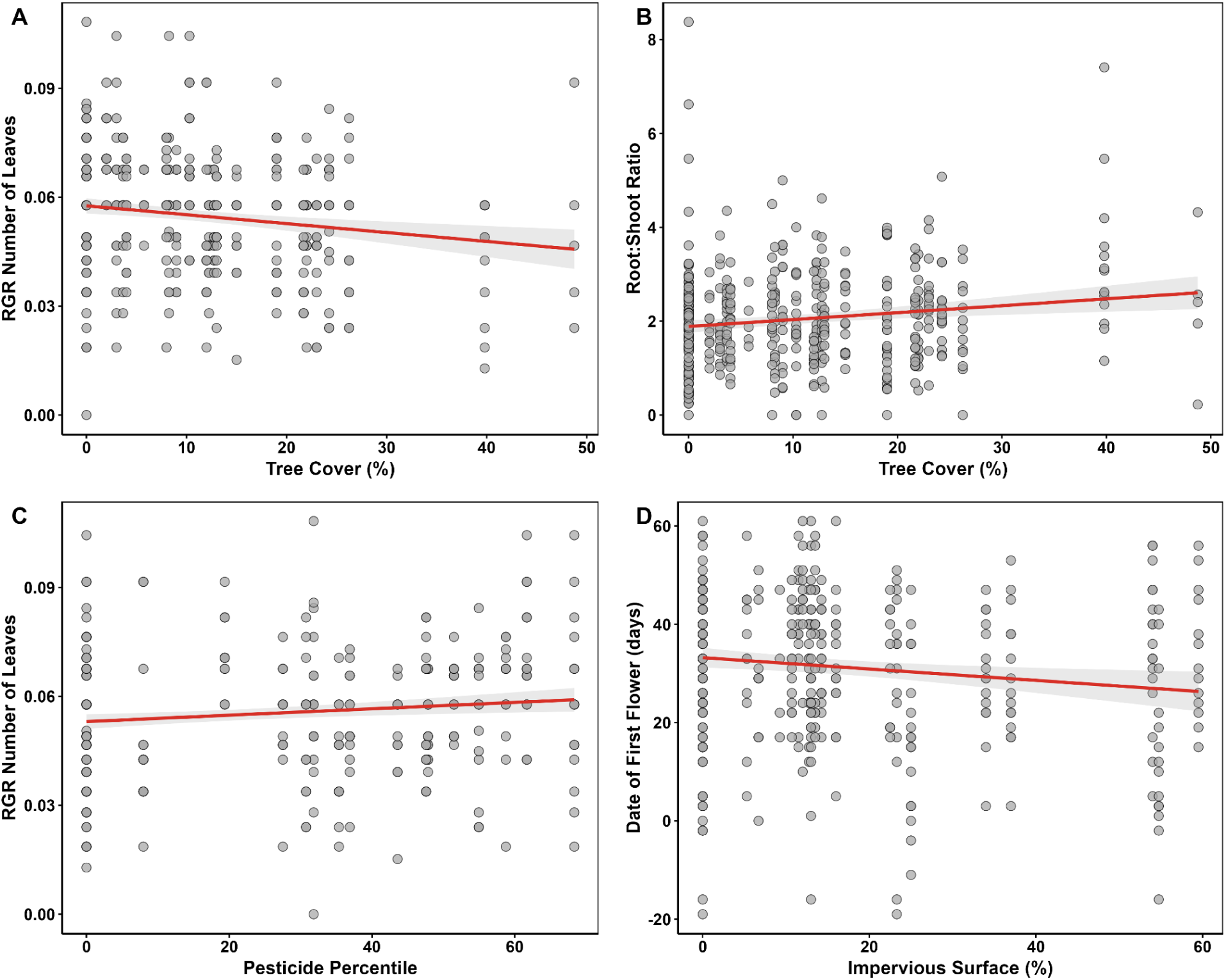
Linear mixed models of the relationship between *Centaurea melitensis* traits and environmental variables. Dots are individual *C. melitensis* plant measurements with a best fit line (red) and 95% confidence interval (gray shading). All relationships were significant prior to Bonferroni corrections, but only the relationship of RGR Number of Leaves with Tree Cover (A) was significant after correction.

Finally, we made an unexpected additional observation that greenhouse block explained an exceptionally high 47.6% of the variance for SLA, while all other traits had less than 20% variance explained by block. Using a linear model to explain SLA by distance from the greenhouse cooling pad, we identified a significant positive relationship (F = 8.287, p = 0.0042) (Figure S3, Table S4). SLA increased with distance from the cooling pad, suggesting that SLA was particularly plastic in our experiment and plants developed thinner leaves as a plastic response to warmer microclimates farther from the cooler.

## DISCUSSION

This analysis leveraged a spatial classification of complex urbanized environments to look for trait differentiation across the urban landscape mosaic, and to test whether trait differences followed the Plant Economic Spectrum predictions for trait-environment strategies in plants (Figure 1). We found that growth traits of *Centaurea melitensis* differed according to Environmental Zones, and were correlated with specific environmental variables associated with urbanization. Though traits demonstrated genetically-based differences across this landscape, we found little evidence for trait variation or differentiation according to the trait-environment strategies predicted by the PES.

### Trait variation

Trait variation across our samples of *C. melitensis* was not indicative of the fundamental trade-offs among trait investments that are predicted by the PES (Figure 1). If such a trade-off had been present, we would have seen negative correlations between suites of traits associated with the resource acquisitive vs. conservative strategies. Instead, the main axis of trait variation (PC1) and correlation matrix indicated a positive correlation among nearly all trait values, reflecting variation from smaller to larger and more fecund individuals across all traits. We did observe negative correlations of varying strengths between the date of first flower and several other variables, such that individuals that flowered earlier also tended to grow faster and produce more flowers and seeds. This is suggestive of a weak axis of variation from ‘slower’ later flowering individuals to ‘faster’ earlier flowering individuals with potentially more resource-acquisitive traits. However, root and shoot investments were positively correlated rather than showing a trade-off between strategies as predicted (Figure 1).

### Trait differentiation among environments

Given that we found no strong axes of multi-trait variation, we tested for differentiation of traits across our urbanized landscape individually for each trait. We found that some traits differed among our multivariate Environmental Zones. In particular, the relative growth rate of both leaf length and the number of leaves were higher in Environmental Zone 4, than nearly all other Zones where we sampled *C. melitensis*. Environmental Zone 4 is dominated by grassland, shrubland, crop, and pastureland, with a high level of pesticide use (Figure S1). Importantly, Environmental Zone 4 and our collections within this Zone are distributed throughout the state (Figure 2), demonstrating the explanatory power of identifying similar habitats in multivariate environmental space for analyses of trait differentiation across landscapes.

Relative to other Zones occupied by *C. melitensis*, Environmental Zone 4 is particularly high in pesticide use, and particularly low in impervious surface and tree cover (Dant et al. 2025). We predicted how each of these variables aligned with the PES resource-acquisitive to resource-conservative axis. We then examined whether trait relationships with these variables followed the associated predictions, and found mixed results. At least one trait showed a trend with each of these variables, but only one of these (the rate of production of leaves versus tree cover) was significant after Bonferroni correction for multiple tests. Given that any local adaptation in an invader is evolutionarily recent and could be weak in magnitude, we discuss each of these trends for its potential ecological implications, but we emphasize that statistical support is weak for all but the relationship of growth to tree cover.

For pesticide use, we predicted a more resource-acquisitive strategy in areas of high pesticide use relative to areas with lower use. This is because pesticides are associated with agricultural environments in this Zone, which are typically thought to provide abundant but ephemeral resources for weedy species such as *C. melitensis* (Petit et al., 2011; Chauhan et al., 2012). Pesticides are also used across many non-agricultural habitats, including the use of herbicide in high quantities in dense urban areas to control invasive and unwanted plants (Sandstrom et al., 2022; Parven et al., 2024). The rapid evolution of herbicide resistance in invasive plants is well-known (Michel et al., 2004; Baucom, 2019; Damalas & Koutroubas, 2024), but other types of trait evolution in response to pesticides across urbanized environments has received little attention to date. We found only one trait correlated with pesticide use, which was the aboveground growth rate as measured by the rate of production of new leaves. RGR of number of leaves was positively related to pesticide use, consistent with a more resource-acquisitive strategy in areas of high pesticide use, but no other traits were differentiated according to pesticide use.

Impervious surface is the most common metric of urbanization (Ramalho & Hobbs, 2012; Szulkin et al., 2020; Murray-Stoker et al., 2025). We predicted a more resource-acquisitive strategy in areas of high impervious surface areas relative to areas with lower impervious surface cover, due to shorter growing seasons in urban heat islands. We found that *C. melitensis* genotypes from sites with greater impervious surface flowered significantly earlier, consistent with a more resource-acquisitive strategy for this trait. Earlier phenology within urban environments has been observed in several previous studies of plants (Jeong et al., 2011; Li et al., 2019; Büntgen et al., 2022; Fuccillo Battle et al., 2022), and is thought to be favored due to shorter growing seasons under the higher temperatures created by the urban heat island effect (Johnson et al., 2015; Čeplová et al., 2017; Woudstra et al., 2024). Previous work has also found that plants in heavily urbanized areas have greater aboveground growth than those in more natural areas (Lambrecht et al., 2016; Yakub & Tiffin, 2017; Santangelo et al., 2020; Qu et al., 2022), which could be consistent with an acquisitive strategy for those species, but we do not see this relationship in *C. melitensis*. Instead, aboveground growth traits were greater in genotypes from grassland and agricultural areas (Environmental Zone 4), and growth traits had no divergence according to the cover of impervious surface.

Finally, for tree cover, we predicted a more resource-acquisitive strategy where tree cover is higher. This is because tree cover in California increases with higher water availability, and it is made up of a combination of forested areas and suburban human landscaping in areas that would otherwise be grass or shrublands (Clarke et al., 2013; Avolio et al., 2015). Higher tree cover might also imply a lower light environment, which would also be expected to select for greater aboveground (more resource acquisitive) investment (Reich, 2014). In contrast to these predictions, we found that genotypes from higher tree cover sites had both slower growth of leaf numbers and higher investment in roots, both of which are more consistent with a resource-conservative strategy (opposite our predictions), though we found no other trait correlations with tree cover. Importantly, greater tree cover at the census tract scale does not mean that *C. melitensis* is growing beneath trees at these sites, and we generally found it in the open when collecting (A.J.D. pers. obs.). Weedy sites with *C. melitensis* in these areas might be relatively nutrient poor, in comparison to agricultural and grassland areas, explaining its increased investment in roots (McNaughton, 1968; Seabloom et al., 2003). In addition, aboveground competition among the herbaceous community could be greater in open grassland and agricultural areas than in the weedy spaces among a higher density of trees, which could also select for greater aboveground investment in regions without trees. Our results suggest that a more specific investigation of environmental variables that differ locally among sites with differing levels of tree cover would be valuable in understanding the environments experienced by this species.

### Implications for understanding adaptation in urban environments

A main finding of our study is that trait variation in this system is not organized along multivariate axes that correspond to the axes of variation proposed by the PES, and that some traits differ across environments in ways that might be predicted by the PES, but that these are not part of predicted multi-trait strategies. We see three potential explanations for the lack of multi-trait PES strategies in this system. First, the PES may not be informative for describing trait differences within annual species (i.e. these strategies do not exist across annual plant populations generally or *C. melitensis* specifically). Second, the PES describes adaptations and the trait differentiation we see may not be adaptive. Third, the trait differentiation we observe may be adaptive and consistent with the PES, but traits may not be evolving at the same rates. We discuss each of these further below.

Whether the PES hypothesis will predict adaptive strategies within annual plant species remains an open question. The PES usually is applied to perennial plant species (de la Riva et al., 2016; Zhao et al., 2017; Volaire, 2018). To our knowledge, there are only two previous studies that have tested its predictions within annual plant species. The first identified traits within *Helianthus anomalus* in the southwestern United States associated with specific environmental stressors (Brouillette et al., 2014). This study found support for trait differentiation consistent with a more resource-acquisitive strategy in more ephemeral environments, and a more resource-conservative strategy in less ephemeral environments, as predicted by the PES. The second study tested for trait associations within 30 winter annuals in Western Asia, and found no significant trait associations as predicted by the PES within any species. The authors questioned the ability of PES to predict trait associations within winter annual species (Kurze et al., 2021). Our study suggests that some traits may follow PES predictions, but that multi-trait strategies may not differentiate among environments as predicted.

An important question remaining from our study is the underlying cause of trait differentiation during this invasion. Trait differentiation could reflect adaptation (what the PES strives to explain), if differentiation is the product of either rapid adaptation during/after invasion or selective establishment of different introductions from different pre-adapted sources. Rapid evolution is well-documented in invasive species, and often appears to be adaptive (Colautti & Lau, 2015; Oduor et al., 2016; Hodgins et al., 2025). Multiple introductions are also common in invasions, and so selective establishment is possible (Dlugosch & Parker, 2008; Uller & Leimu, 2011; Dlugosch et al., 2015). However, for *C. melitensis,* we detected trait differentiation among populations in different Environmental Zones that are geographically distributed and interspersed with other Zones, reducing the likelihood that the trait differentiation can be explained by introductions from different sources into different areas. Trait differentiation may also reflect random differences among environments due to genetic drift or founder effects. We cannot rule out this possibility without further population genetic studies (see Appendix D of this dissertation).

A feature shared by all species invasions (by definition) is that they are evolutionarily recent, and local adaptation is likely to be constrained by the rate of adaptation of traits, especially traits constrained by genetic covariances with other traits (Colautti & Barrett, 2011; Blows & McGuigan, 2015; McGoey & Stinchcombe, 2021). While *C. melitensis* has been in California for over 200 years, this may have been only enough time for some traits to locally adapt. For example, we observed that flowering time is earlier in lineages from sites with higher impervious surface area, but only flowering time was differentiated according to impervious surface area. Flowering time is known to be one of the most rapidly evolving traits in plants, with a clear genetic basis and standing variation of alleles with large effects on the trait (Franks et al., 2007; Zan & Carlborg, 2019; Yan et al., 2021). It has repeatedly been observed to evolve adaptively in response to single drought events (Misson et al., 2011; Tun et al., 2021). In contrast, other traits may be slower to adapt locally due to lower genetic variation or greater constraints.

The trait variation that we quantified does reflect a potential for this invader to adapt to environmental variables across its landscape, including to human modifications of the environment (Moran & Alexander, 2014; Johnson & Munshi-South, 2017; Khattak et al., 2024). Notably, the region with greatest trait differentiation from others occupied by *C. melitensis* was

Environmental Zone 4, which is dominated by grasslands and agricultural lands and is particularly low in impervious surfaces. This suggests that most populations share traits suitable for occupying more heavily urbanized areas, and any adaptation to more natural grasslands and agricultural areas would be more recent in this invasion of *C. melitensis*. The idea that cities and other anthropogenically modified spaces can create opportunities for invasive species evolution into natural areas is one that has been previously discussed by Borden and Flory 2021. They suggested that invasive species may first colonize and adapt to urban areas in their native range, enabling these species to spread to other cities outside of their native range, from which they might invade natural areas in their introduced range. The native habitat from which *C. melitensis* arrived in California is unknown, but the oldest record of *C.melitensis* was found in an adobe brick dated to 1789, suggesting close proximity to human disturbance.

Finally, we note that trait plasticity is also an important component of plant trait strategies and adaptation to the environment, and is likely to play a role in the adaptive strategies of *C. melitensis*. The trait-environment relationships found here differ from those identified in an earlier study of first-generation plants grown from field-collected seeds (and a smaller number of collection sites; Dant et al. 2025), indicating environmental effects. Within this study, we did not quantify plasticity directly but we found a significant block effect (plastic variation) in one trait: specific leaf area (SLA). SLA was significantly affected by the positioning in the greenhouse, such that blocks which were nearer to cooling pads had thicker leaves while those further away from the cooling pads were thinner. Other plastic responses have also been reported within *C. melitensis,* including growth rate plasticity within its invaded North America range, but not in its native, European range, suggesting the evolution of increased plasticity (Moroney et al., 2013). Germination plasticity within invaded populations in Australia has also been observed (Humphries et al., 2024). Studies that quantify both plastic and genetic variation in traits, and their fitness consequences, would be valuable for further understanding the potential for adaptive responses to urbanization and complex environmental variation in this system.

### Conclusions

Our study identified evidence of genetically-based differentiation of multiple traits in the *C. melitensis* invasion of complex urbanized environments in California, USA. The relationships between these traits and environmental variables were consistent with some predictions of the PES, but in all cases the expected multi-trait strategies were lacking. This suggests that either the PES is not informative for interpreting local adaptation in this annual species, or that its trait divergence is not adaptive, or that traits are adapting independently and at different rates. These results indicate a need for further tests of the adaptive nature of these differences but also highlight the opportunities for urban areas to serve as stepping stones for future adaptation to natural areas in this invasion.

## Open research statement

All raw data and code for this manuscript can be found at https://github.com/adantdudleya/Dant_etal_2026_Cenmel_Greenhouse. The repository includes (1) the raw greenhouse trait data (Master_Data_6_30_2025.csv) and the greenhouse block layout (Block_data.csv), (2) spatial data for 7,829 California census tracts (urban_typology2.shp and associated files), and (3) two R scripts (Chapter2_Trait_Analysis.R and Ch2_California_Maps.R) implementing the analyses described in this study. Upon acceptance, a permanent version of the repository will be archived in Zenodo with a DOI.

## ACKNOWLEDGMENTS

We would like to thank S. King, and T. Reese for assistance in the greenhouse. In addition, we would like to thank J. Autz for helping manage the University of Arizona 8th street greenhouse. Finally, we thank J. Bronstein and G. Barron-Gafford for helpful comments on earlier versions of this manuscript.

## DATA AVAILABILITY

All raw data and code underlying this study are openly available on GitHub at https://github.com/adantdudleya/Dant_etal_2026_Cenmel_Greenhouse. The repository includes (1) the raw greenhouse trait data (Master_Data_6_30_2025.csv) and the greenhouse block layout (Block_data.csv), (2) spatial data for 7,829 California census tracts (urban_typology2.shp and associated files), and (3) two R scripts (Chapter2_Trait_Analysis.R and Ch2_California_Maps.R) implementing the analyses described in this study. Upon acceptance, a permanent version of the repository will be archived in Zenodo with a DOI.

## AUTHOR CONTRIBUTIONS

Anthony Dant conceived the study. Anthony Dant and Katrina Dlugosch designed and secured funding for the research. Anthony Dant and Lily Bishop conducted the research. Anthony Dant conducted analyses and drafted the manuscript, with assistance from Katrina Dlugosch. All authors reviewed and approved the final version of the manuscript.

## AI STATEMENT

The authors used a generative AI assistant to help edit and troubleshoot R code for data analysis and figure generation, and to assist with language editing. The authors reviewed and verified all code and text, and take full responsibility for the content of this manuscript.

