## Appendix S1 for "Trait differentiation across urbanized environments in invasive *Centaurea melitensis* is not predicted by the Plant Economic Spectrum"

Title: Trait differentiation across urbanized environments in invasive *Centaurea melitensis* is not predicted by the Plant Economic Spectrum

Authors: Anthony J. Dant, Lily G. Bishop, Katrina M. Dlugosch

**SUPPLEMENTARY TABLES**

**Table S1.** Collection site metadata for 31 *Centaurea melitensis* populations sampled across California.

| **Site** | **Environmental Zone** | **Region** | **Latitude** | **Longitude** | **N (individuals)** | **Tree Cover (%)** | **Impervious Surface (%)** | **Pesticide Percentile** |
| --- | --- | --- | --- | --- | --- | --- | --- | --- |
| Balbon | 1 | Southern | 32.7381 | -117.1428 | 17 | 9 | 13 | 0 |
| Torres | 1 | Southern | 32.8883 | -117.2492 | 17 | 4 | 16 | 0 |
| University | 1 | Southern | 32.7753 | -117.1911 | 16 | 0 | 0 | 0 |
| Cal Poly | 3 | Central | 35.3089 | -120.6547 | 18 | 24.25 | 0 | 54.97 |
| Dark | 3 | Southern | 34.4647 | -119.6881 | 14 | 22 | 0 | 27.52 |
| Garapata | 3 | Central | 36.4544 | -121.9242 | 15 | 23 | 11.5 | 0 |
| Garden | 3 | Northern | 37.0497 | -122.0831 | 14 | 19 | 22.5 | 0 |
| Lobos | 3 | Northern | 37.0497 | -122.0831 | 5 | 48.75 | 9.25 | 0 |
| Long | 3 | Northern | 37.1594 | -121.8758 | 14 | 26.25 | 12 | 30.76 |
| Marco | 3 | Southern | 34.4567 | -119.7667 | 14 | 0 | 5.3 | 47.59 |
| Oro | 3 | Central | 35.2581 | -120.8872 | 15 | 8 | 0 | 58.76 |
| Point Reyes | 3 | Northern | 38.0028 | -122.8122 | 14 | 39.8 | 0 | 0 |
| Seaside | 3 | Central | 36.6439 | -121.8053 | 17 | 0 | 59.5 | 0 |
| Pina | 4 | Central | 36.4814 | -121.1817 | 17 | 10.3 | 0 | 61.69 |
| Ghost | 5 | Southern | 33.54 | -117.5564 | 18 | 12.75 | 14.25 | 47.86 |
| King | 5 | Southern | 34.1022 | -118.7092 | 14 | 12 | 0 | 7.97 |
| Old | 5 | Southern | 32.8394 | -117.0414 | 15 | 8.25 | 13 | 0 |
| Ranch | 5 | Southern | 34.1083 | -118.7492 | 12 | 2 | 0 | 19.38 |
| Sand | 5 | Southern | 34.1072 | -118.9092 | 16 | 15 | 0 | 43.59 |
| Tuna | 5 | Southern | 34.0183 | -119.6822 | 6 | 5.75 | 0 | 0 |
| Border | 6 | Southern | 32.5447 | -117.0742 | 16 | 0 | 6.7 | 35.38 |
| Don | 6 | Northern | 37.4356 | -121.9581 | 12 | 0 | 12.75 | 0 |
| Irish | 6 | Central | 35.2514 | -120.6933 | 16 | 3 | 13.5 | 68.34 |
| Mariana | 6 | Southern | 33.9656 | -118.4269 | 18 | 21.7 | 37 | 0 |
| San Diego | 6 | Southern | 32.6394 | -117.1017 | 18 | 3.66 | 54 | 0 |
| San Elijo | 6 | Southern | 33.0133 | -117.2736 | 14 | 13 | 54.74 | 36.9 |
| CSU Dom | 8 | Southern | 33.8589 | -118.255 | 18 | 0 | 25 | 31.83 |
| Debb | 8 | Southern | 34.1203 | -118.2869 | 15 | 19 | 23.3 | 0 |
| Franklin | 8 | Southern | 34.1236 | -118.4089 | 14 | 12.25 | 0 | 0 |
| Griff | 8 | Southern | 34.0994 | -118.2008 | 17 | 0 | 34 | 0 |
| Lake | 9 | Southern | 34.1836 | -118.4997 | 17 | 0 | 10.7 | 51.48 |
| **Total** |  |  |  |  | **463** |  |  |  |

*Sites are sorted by Environmental Zone. Environmental variables (tree cover, impervious surface, pesticide percentile) were extracted at the census tract level as described in the methods. N = number of individuals with trait data in the common garden experiment.*

**Table S2.** Linear mixed model results for functional traits predicted by Environmental Zone with Site nested within Zone. An asterix denotes significant *P*-values after Bonferroni correction.

| **Trait** | **N** | **Zone F** | **Zone p** | **Site F** | **Site p** | **Zone 1** | **Zone 3** | **Zone 4** | **Zone 5** | **Zone 6** | **Zone 8** | **Zone 9** |
| --- | --- | --- | --- | --- | --- | --- | --- | --- | --- | --- | --- | --- |
| RGR Length | 443 | 5.28 | < 0.001* | 2.86 | < 0.001 | a | a | b | ab | a | a | ab |
| RGR Width | 434 | 3.07 | 0.006 | 1.901 | 0.007 | a | a | a | a | a | a | a |
| RGR Number of Leaves | 451 | 3.76 | 0.001* | 3.096 | < 0.001 | a | a | b | a | a | a | ab |
| Specific Leaf Area (cm²/g) | 447 | 0.76 | 0.602 | 1.844 | 0.0096 | — | — | — | — | — | — | — |
| Aboveground Biomass (g) | 448 | 1.57 | 0.154 | 1.284 | 0.1683 | — | — | — | — | — | — | — |
| Root:Shoot Ratio | 440 | 1.06 | 0.385 | 2.484 | 0.0002 | — | — | — | — | — | — | — |
| Date of First Flower (days) | 393 | 3.53 | 0.002* | 2.683 | < 0.001 | ab | b | ab | ab | a | ab | ab |
| Number of Flowers | 430 | 1.11 | 0.356 | 3.297 | < 0.001 | — | — | — | — | — | — | — |
| Average Seed Weight (g) | 388 | 1.67 | 0.128 | 1.151 | 0.287 | — | — | — | — | — | — | — |

*Post-hoc groups: Sidak-adjusted pairwise comparisons via emmeans. Zones sharing the same letter are not significantly different (α = 0.05). Highlighted cells (b) indicate zones significantly different from the majority group (a). Dashes indicate a Zone effect was not significant.*

**Table S3.** Linear mixed model results for functional traits predicted by source population environmental variables. Bolded values are significant. An asterix denotes significant *P*-values after Bonferroni correction. Variance inflation factors (VIF) showed there was no collinearity among any of the three environmental variables (VIF range: 1.07–1.16).

| **Trait** | **Environmental Variable** | **N** | **Estimate** | **Std. Error** | **t** | **F** | **p** |
| --- | --- | --- | --- | --- | --- | --- | --- |
| RGR Length | Impervious Surface (%) | 443 | -0.00003 | 0.00007 | -0.416 | 0.173 | 0.6778 |
| RGR Length | Pesticide Percentile | 443 | 0.00001 | 0.00005 | 0.171 | 0.029 | 0.8645 |
| RGR Length | Tree Cover (%) | 443 | -0.00013 | 0.0001 | -1.254 | 1.572 | 0.2106 |
| RGR Width | Impervious Surface (%) | 434 | -0.00005 | 0.00004 | -1.554 | 2.416 | 0.1208 |
| RGR Width | Pesticide Percentile | 434 | -0.00001 | 0.00002 | -0.454 | 0.206 | 0.6504 |
| RGR Width | Tree Cover (%) | 434 | 0 | 0.00005 | -0.046 | 0.002 | 0.963 |
| RGR Number of Leaves | Impervious Surface (%) | 451 | -0.00001 | 0.00005 | -0.296 | 0.088 | 0.7671 |
| RGR Number of Leaves | Pesticide Percentile | 451 | 0.00007 | 0.00003 | 2.033 | **4.133** | **0.0426** |
| RGR Number of Leaves | Tree Cover (%) | 451 | -0.00021 | 0.00007 | -3.054 | **9.325** | **0.0024*** |
| Specific Leaf Area (cm²/g) | Impervious Surface (%) | 447 | 0.16907 | 0.23329 | 0.725 | 0.525 | 0.469 |
| Specific Leaf Area (cm²/g) | Pesticide Percentile | 447 | 0.07258 | 0.16548 | 0.439 | 0.192 | 0.6611 |
| Specific Leaf Area (cm²/g) | Tree Cover (%) | 447 | 0.59505 | 0.35277 | 1.687 | 2.845 | 0.0924 |
| Aboveground Biomass (g) | Impervious Surface (%) | 448 | 0.00021 | 0.00013 | 1.554 | 2.413 | 0.121 |
| Aboveground Biomass (g) | Pesticide Percentile | 448 | 0.00011 | 0.00009 | 1.12 | 1.254 | 0.2634 |
| Aboveground Biomass (g) | Tree Cover (%) | 448 | -0.00013 | 0.0002 | -0.624 | 0.39 | 0.5328 |
| Root:Shoot Ratio | Impervious Surface (%) | 440 | -0.00332 | 0.00309 | -1.076 | 1.158 | 0.2825 |
| Root:Shoot Ratio | Pesticide Percentile | 440 | -0.0008 | 0.0022 | -0.366 | 0.134 | 0.7149 |
| Root:Shoot Ratio | Tree Cover (%) | 440 | 0.01104 | 0.00471 | 2.342 | **5.486** | **0.0196** |
| Date of First Flower (days) | Impervious Surface (%) | 393 | -0.09988 | 0.04624 | -2.16 | **4.666** | **0.0314** |
| Date of First Flower (days) | Pesticide Percentile | 393 | -0.0343 | 0.03232 | -1.061 | 1.126 | 0.2893 |
| Date of First Flower (days) | Tree Cover (%) | 393 | 0.05021 | 0.07241 | 0.693 | 0.481 | 0.4885 |
| Number of Flowers | Impervious Surface (%) | 430 | 0.00472 | 0.0043 | 1.098 | 1.206 | 0.2727 |
| Number of Flowers | Pesticide Percentile | 430 | 0.00296 | 0.00306 | 0.967 | 0.935 | 0.3341 |
| Number of Flowers | Tree Cover (%) | 430 | -0.01219 | 0.00662 | -1.843 | 3.396 | 0.0661 |
| Average Seed Weight (g) | Impervious Surface (%) | 388 | 0.00308 | 0.00308 | 1.00 | 1.001 | 0.3178 |
| Average Seed Weight (g) | Pesticide Percentile | 388 | 0.00096 | 0.00216 | 0.446 | 0.199 | 0.6561 |
| Average Seed Weight (g) | Tree Cover (%) | 388 | 0.00052 | 0.00481 | 0.108 | 0.012 | 0.9137 |

**Table S4.** Variance components for greenhouse block position from environmental variable linear mixed models.

| **Trait** | **Block Variance** | **Residual Variance** | **Total Variance** | **Block (%)** |
| --- | --- | --- | --- | --- |
| RGR Length | 7.37E-05 | 4.69E-04 | 5.42E-04 | 13.59 |
| RGR Width | 1.61E-05 | 1.30E-04 | 1.46E-04 | 11.02 |
| RGR Number of Leaves | 3.94E-05 | 2.24E-04 | 2.63E-04 | 14.97 |
| **Specific Leaf Area (cm²/g)** | **5338.871** | **5884.341** | **11223.212** | **47.57** |
| Aboveground Biomass (g) | 4.57E-04 | 1.95E-03 | 2.41E-03 | 18.96 |
| Root:Shoot Ratio | 0.027 | 1.038 | 1.064 | 2.52 |
| Date of First Flower (days) | 33.647 | 207.012 | 240.659 | 13.98 |
| Number of Flowers | 0.174 | 1.963 | 2.137 | 8.13 |
| Average Seed Weight (g) | 0.0124 | 0.903 | 0.916 | 1.36 |

*Block variance represents the proportion of total variance attributable to block position in the greenhouse (proximity to evaporative cooling pad). The SLA row is highlighted due to its high block variance (47.6%), which is explored further in the supplementary figures.*

**SUPPLEMENTARY FIGURES**

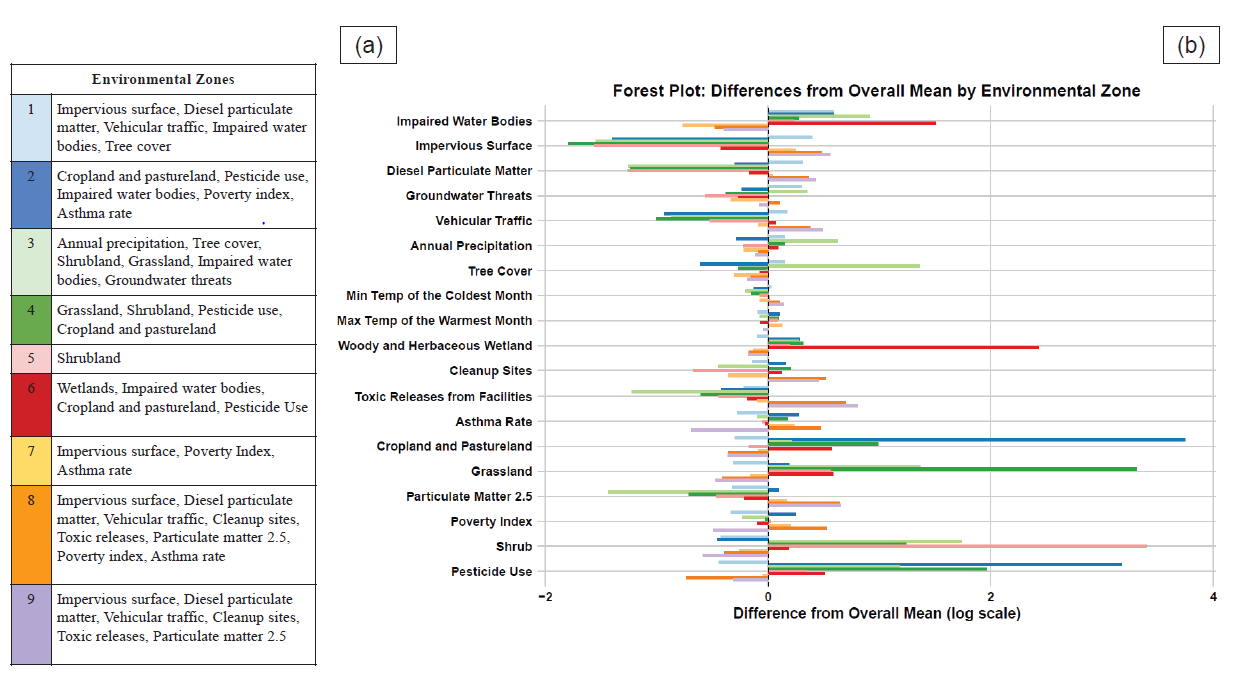

**Figure S1.** Forest plot of the mean values of all 19 environmental variables used in the hierarchical cluster analysis, for each of the nine Environmental Zones (colors) described in Dant et al. 2025. (a) Legend listing the environmental characteristics most associated with each of the nine Environmental Zones. (b) Forest plot showing differences in mean log-transformed values of the 19 environmental variables for each Environmental Zone. The x-axis represents the difference between the mean of log(x+1) for a variable within a specific Environmental Zone and the overall mean of log(x+1) across all 7,829 census tracts. See Dant et al. 2025 for further details.

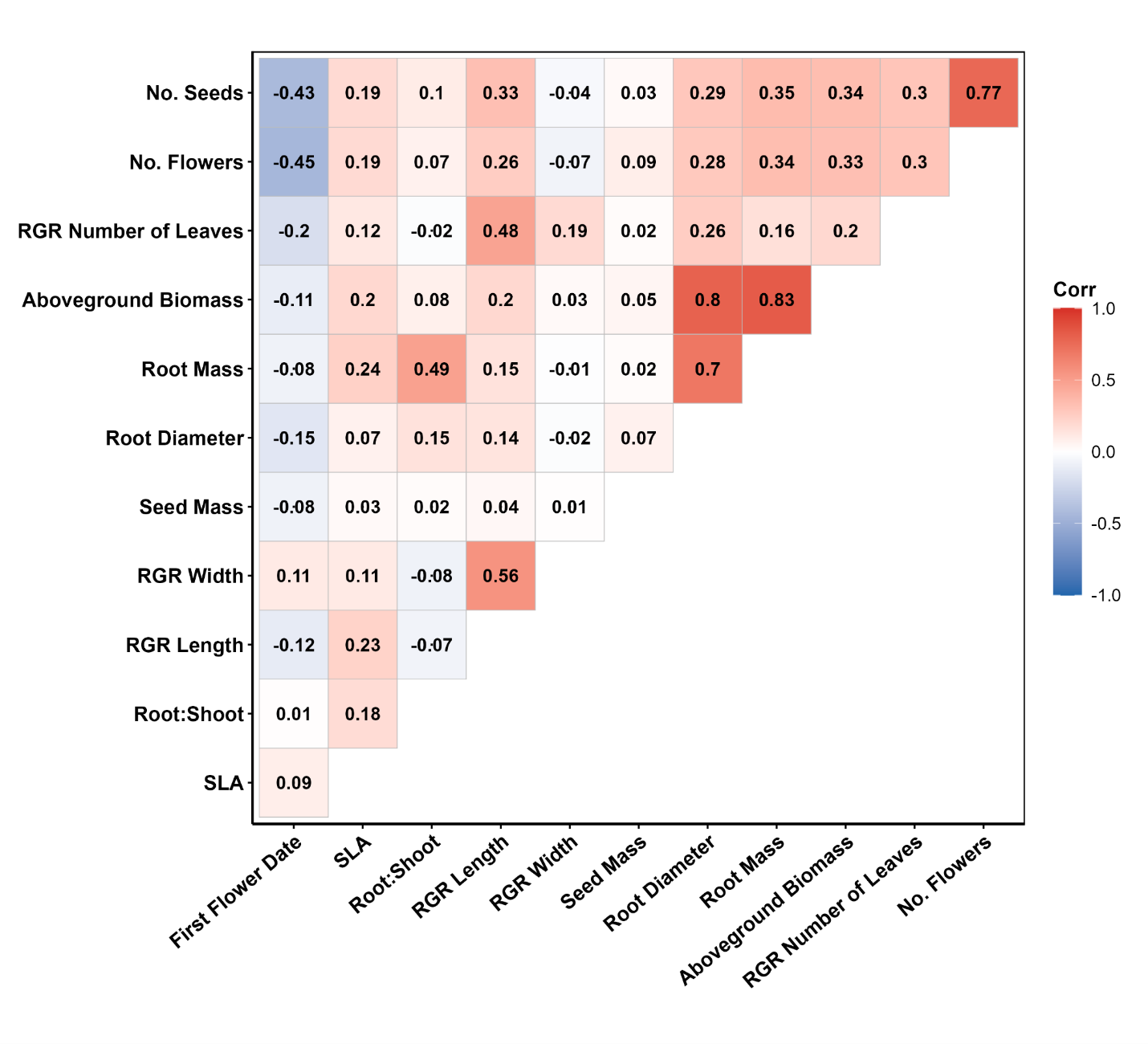
**Figure S2.** Correlation matrix of all 12 traits colored by level of correlation and direction of relationship (red positive and blue negative). Root diameter and root mass were excluded in further analyses in favor of above ground biomass. The total number of seeds was excluded in favor of the total number of flowerheads.

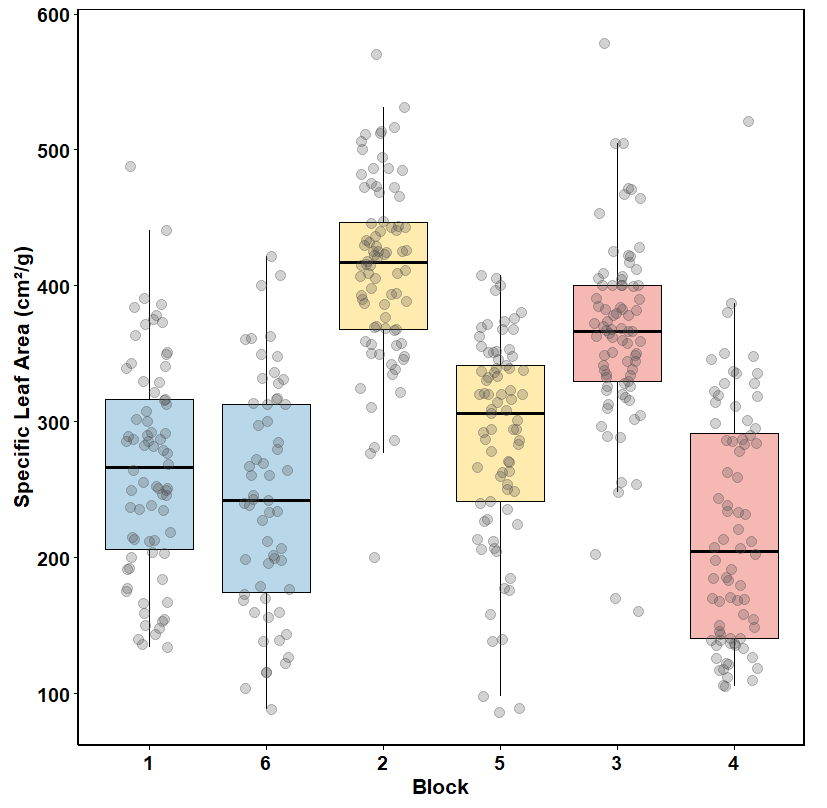
**Figure S3.** Association between Specific Leaf Area (cm2/g) measured on *C. melitensis* individuals and position within the greenhouse. Blocks are color coded and grouped according to their position relative to cooling pads within the greenhouse with Blocks 1 and 6 being the closest (blue), and 3 and 4 being the furthest away (red).
